# Ecological genetic conflict between specialism and plasticity through genomic islands of divergence

**DOI:** 10.1101/298554

**Authors:** Olof Leimar, Sasha R. X. Dall, John M. McNamara, Bram Kuijper, Peter Hammerstein

## Abstract

There can be genetic conflict between genome elements differing in transmission patterns, and thus in evolutionary interests. We show here that the concept of genetic conflict provides new insight into local adaptation and phenotypic plasticity. Local adaptation to heterogeneous habitats sometimes occurs as tightly linked clusters of genes with among-habitat polymorphism, referred to as genomic islands of divergence, and our work sheds light on their evolution. Phenotypic plasticity can also influence the divergence between ecotypes, through developmental responses to habitat-specific cues. We show that clustered genes coding for ecological specialism and unlinked generalist genes coding for phenotypic plasticity differ in their evolutionary interest. This is an ecological genetic conflict, operating between habitat specialism and phenotypically plastic generalism. The phenomenon occurs both for single traits and for syndromes of co-adapted traits. Using individual-based simulations and numerical analysis, we investigate how among-habitat genetic polymorphism and phenotypic plasticity depend on genetic architecture. We show that for plasticity genes that are unlinked to a genomic island of divergence, the slope of a reaction norm will be steeper in comparison with the slope favored by plasticity genes that are tightly linked to genes for local adaptation.

## Introduction

Genetic conflict occurs when different genomic elements, or different haplotypes at a locus, differ in their evolutionary interests. This possibility has been given much attention (Hurst et al. 1996; Werren and Beukeboom 1998; Burt and Trivers 2006; Gardner and Úbeda 2017), resulting in the insight that genetic conflict can be important for evolutionary change and innovation, as well as influence phenomena like sex determination (Werren 2011). Most work has focused on genetic conflict with a basis in the properties of genetic transmission systems. Thus, different pathways of transmission, as for nuclear and mitochondrial genes (e.g., Frank and Hurst 1996; Perlman et al. 2015), or the biasing of transmission along a pathway, as for segregation distorters (Hurst et al. 1996), have been put forward as sources of genetic conflict. Ecologically based genetic conflict is, however, a possibility that is less well-established in evolutionary biology. In heterogeneous environments, genes can differ in their pathways of transmission to future generations, involving the kinds of environments they pass through. Genetic conflict can then appear by favoring or suppressing these pathways, and thus have a basis in the ecology of populations (Leimar et al. 2006; Dall et al. 2015; Leimar et al. 2016).

The long-term reproductive success of a gene, in the sense of its representation in future generations, defines its evolutionary interest. For heterogeneous habitats, the question arises in which kind of habitat we should count future representation. For genes that code for a specialist phenotype it is the long-term representation in the habitat for which the phenotype is specialized that defines evolutionary interest, but for genes for generalism it is instead the representation over a range of habitats that matters. This general idea, with a focus on two habitats and a phenotypically plastic generalist, is illustrated in fig. 1. Although genes for specialism may, as a result of dispersal, end up in different habitats, they are selected against in habitats to which they are not adapted, and have little evolutionary future there. This is an example of a source-sink process (Holt and Gaines 1992; Kawecki 1995), entailing that a specialist already adapted to one habitat need not evolve to be adapted to another, even if there is a certain amount of migration between habitats. Concerning plasticity, we assume that it is a developmental response to a noisy environmental cue.

**Figure 1:**
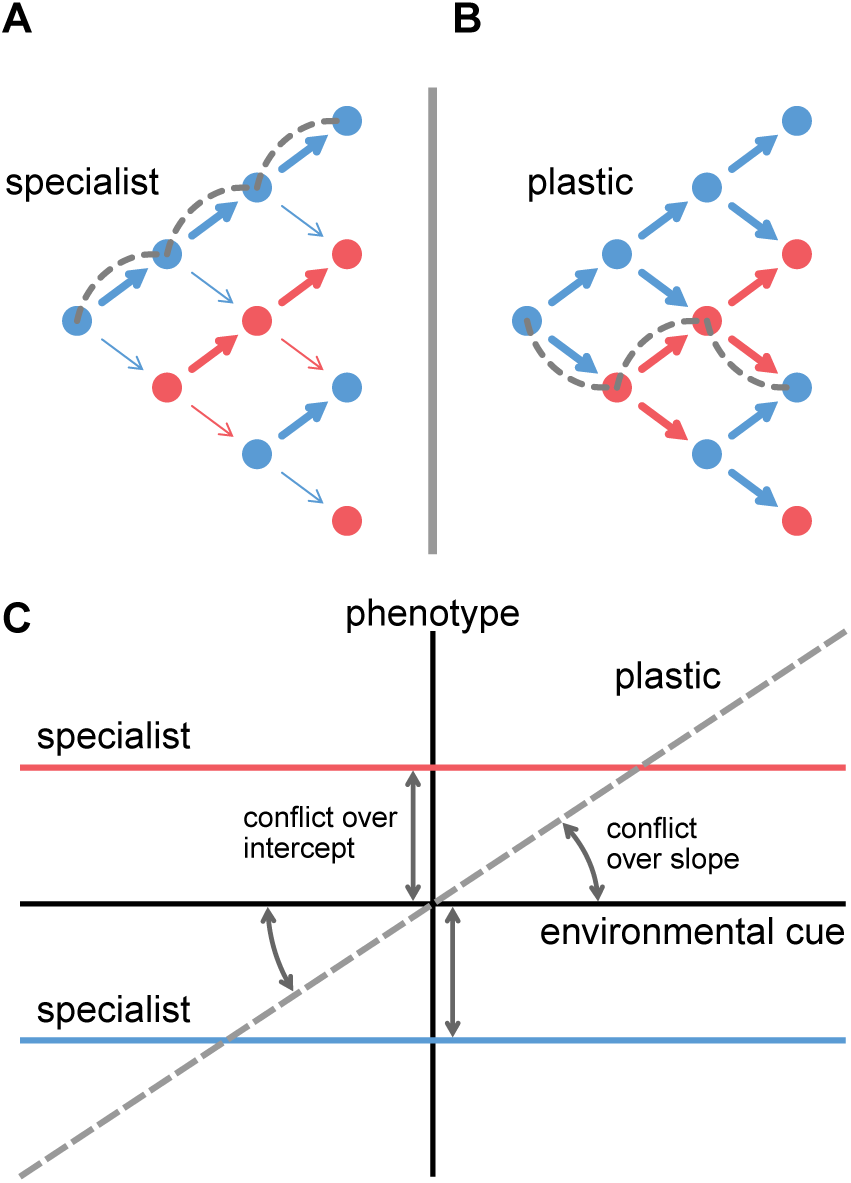
An overview of ecological genetic conflict between genes for specialism and for phenotypically plastic generalism. Illustrations of different pathways of transmission to future generations for *A* genes for habitat specialism and *B* plasticity, and *C* a resulting conflict battleground. Blue and red indicate two different habitats and the arrows show potential dispersal events, when changes between habitat types are possible. Time runs from left to right in panels *A* and *B*. For a specialist *A*, a pathway of transmission to future generations will predominantly go trough one habitat type, illustrated by the dashed gray line, because the alternative habitat is a sink. For a phenotypically plastic generalist *B*, on the other hand, a pathway of transmission to future generations can alternate between habitat types (e.g., dashed gray line). As a consequence, there will be a divergence in evolutionary interests between genes for specialism and plasticity. *C* Locally adapted genes (specialist) are then in conflict with genes for plasticity (generalist) over both the intercept and the slope of reaction norms.

The difference in evolutionary interests illustrated in fig. 1A,B corresponds to the general idea of genetic conflict (Gardner and Úbeda 2017), and it is thus appropriate to refer to it as an ecological genetic conflict. In analyzing genetic conflict, it is essential to take into account which traits are influenced by genes that differ in their evolutionary interest. Since we study genetic local adaptation and plasticity, we focus on intercepts and slopes of reaction norms (fig. 1C). For instance, pure local adaptation in heterogeneous habitats could be implemented as a genetically polymorphic locus influencing the intercepts in fig. 1C, with slopes being flat. A pure phenotypically plastic generalist, on the other hand, would be implemented as genetically monomorphic intercept and slope. Often, the evolutionary outcome is intermediate between these. Our aim is to explain the role ecological genetic conflict plays in influencing the outcome.

Genetic architecture is an important aspect of our analysis, in particular how genes influencing the intercept and slope of a reaction norm (fig. 1C) are positioned in the genome. To see why linkage matters, note that if two genes are fully linked, their evolutionary interests must coincide, because their representation in a future gene pool will be the same. As a consequence, only genes with less than full linkage, e.g. unlinked genes, can differ in their evolutionary interest. The latter includes genes in the same position, but on different physical stretches of DNA, such as different genes at a locus. Two genes at the same locus but locally adapted to different habitats can differ in their evolutionary interest, together with genes tightly linked to each of them.

It is a characteristic feature of genetic conflict that the degree of linkage between genes influences the evolutionary outcome. There is a tendency for a segment of DNA to share the evolutionary interest of selfish elements (Hurst et al. 1996; Burt and Trivers 2006), which can result in the formation of ‘selfish’ co-adapted gene complexes or supergenes (for definitions of these terms, see Schwander et al. 2014). As we show here, a similar principle applies to ecological genetic conflicts, where a genomic island of divergence (Nosil et al. 2009) can correspond to the shared interest of a number of genes for local adaptation to specific habitats, in this way favoring transmission through certain habitats but not others. The region might contain several genes of smaller effect that add up to a bigger effect for a particular trait (Yeaman and Whitlock 2011; Yeaman 2013). More generally, it can contain genes that epistatically modify the effect of other genes in the region, as well as genes that influence different traits that contribute to local adaptation (Feder et al. 2012; Marques et al. 2016; Larson et al. 2017), making up a co-adapted gene complex. Although the phenomenon of genomic islands of divergence has attracted much interest in recent years (Nosil et al. 2009; Jones et al. 2012; Via 2012; Flaxman et al. 2014; Lucek et al. 2014; Poelstra et al. 2014; Seehausen et al. 2014; Soria-Carrasco et al. 2014; Malinsky et al. 2015; Riesch et al. 2017), the idea of differences in evolutionary interest between genes in an island of divergence and unlinked genes has not been explored, neither has the idea that plasticity genes can favor different reaction norm slopes depending on their linkage to an island of divergence.

Another characteristic feature of genetic conflict is that natural selection will not in general lead to a unique outcome for traits that are influenced by genes in conflict. Instead, there might be an ‘arms race’ (Hurst et al. 1996; Werren 2011), where the outcome is influenced by such things as supply of mutations, position in the genome, and limits to gene expression. Our main way of dealing with this is to examine situations where there is selectively maintained genetic polymorphism at one or more loci, but where we do not focus on the possible evolution of the corresponding genes. Instead, we examine the evolution of genes that modify the phenotypic effects of the polymorphism, for instance modify intercepts and slopes of a reaction norm (fig. 1C). By examining situations where modifiers of intercepts and plasticity genes influencing slopes have tight versus loose linkage to a polymorphic locus, we can examine the effect of genetic architecture. The genetic conflict we investigate is thus between loci that are tightly versus loosely linked to a genetic polymorphism. As we will show, this can be interpreted in terms of a conflict between habitat specialism, corresponding to tight linkage, and phenotypically plastic generalism, corresponding to loose linkage.

We examine between-habitat genetic polymorphism for one trait, as well as for two different traits, for which the optimum differs between habitats, and we determine how the relative contributions of between-habitat genetic polymorphism and phenotypic plasticity depend on genetic architecture. As mentioned, the kind of genetic architecture we are concerned with is the degree of linkage between genetically polymorphic loci, epistatic modifiers of the effects at these loci, and plasticity genes influencing a reaction norm slope. We emphasize the distinction between the case where all loci are tightly linked together in a supergene and that where modifier and plasticity loci are unlinked to genetically polymorphic loci. However, we also investigate intermediate cases, for instance a polymorphic locus with a tightly linked modifier and an unlinked plasticity locus determining the slope of a reaction norm. In such a case, intercept and slope of a reaction norm (fig. 1C) are determined by genes with diverging evolutionary interests.

Among our reasons for examining the combination of genetic differentiation and phenotypic plasticity are, first, that traits of ecotypes in nature often are the combined result of genetic and environmental effects and, second, that comparison of the relative weights in phenotype determination on different inputs, such as genetic polymorphism and environmental cues, gives a striking picture of the influence of genetic architecture on the evolutionary outcome. To make contact with previous work on genomic islands of divergence, we briefly examine the role of linkage between many loci of small effect in building up a larger effect of between habitat genetic polymorphism. In addition to genetic architecture, we examine the influence of the rate of migration between habitats and the strength of selection on the characteristics of local adaptation. We also study the question of the evolution of the rate of recombination between polymorphic loci, modifiers, and plasticity loci. For the analysis, we use individual-based evolutionary simulations of diploid populations, with several local populations in each habitat, as well as numerical analysis of evolutionary equilibria for a model with a very large population in each habitat. For simplicity, we let the sex of an individual be randomly determined (Perrin 2016).

A main finding is that for plasticity genes that are unlinked to a genomic island of divergence, the slope of a reaction norm will be steeper in comparison with the slope favored by plasticity genes that are tightly linked to genes for local adaptation. This holds in particular for intermediate rates of between-habitat migration. We discuss our results in relation to empirical work on the genomics of ecotypic variation and on the relative importance of genetic variation and plasticity for local adaptation.

## Methods

We first present our two-habitat metapopulation model for a single trait *u*, then extend it to two traits *u*_1_ and *u*_2_, followed by an explanation of our individual-based simulations. We have also performed a numerical analysis of a model with a very large population in each habitat, which is described in the supporting information, with results reported in Table A1 and fig. A1.

### Single trait

The population is divided into *N*_*p*_ patches, each containing a local population with on average *K* diploid individuals with non-overlapping generations, and with survival selection operating in each patch. An individual’s sex is randomly determined, and each offspring is formed by randomly selecting a mother and a father from the local population. There is a genotype-cue-phenotype mapping, determining an individual’s phenotype *u* as a weighted sum of a ‘genetic effect’ *z* and a environmental cue *x*_juv_, such that

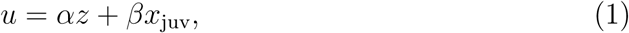

where *z* and the weights *α* and *β* are each determined by a diploid locus. This means that there is epistasis between the locus for the genetic effect *z* and the locus coding for *α*.

A patch is in either of two environmental states, corresponding to two types of habitat, which could, for instance, be low and high resource availability, risk of predation, or salinity. The two habitats are denoted by *i* = 1, 2, with juvenile-to-adult survival for phenotype *u* in habitat *i* given by

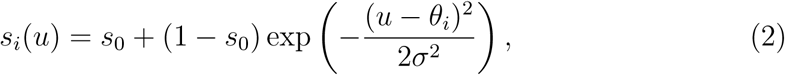

where *s*_0_ is a basic survival rate, *θ*_*i*_ is the optimal phenotype in habitat *i* and *σ* is the width of the Gaussian survival function. An individual can get information about which habitat it is in through the juvenile cue, given by

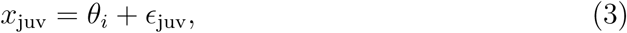

where *θ*_*i*_ is the mean cue in habitat *i*, for simplicity assumed to be the same as the optimal phenotype, and *E*_juv_ is a normally distributed random error with mean 0 and standard deviation *σ*_juv_.

There is a probability *m* of juvenile dispersal to a patch randomly selected in the entire metapopulation, including the patch of origin. The local populations are regulated such that a patch produces *K* juveniles, each of which has a probability *m* to disperse. There are equal numbers of patches for the two habitats, which means that the probability for a dispersing individual to change habitat is *m/*2.

The life cycle of individuals is as follows: (*i*) selection, with survival in habitat *i* as a function of phenotype *u* as in equation (2); (*ii*) within-patch random mating, forming *K* offspring in each local population, after which the adults die; (*iii*) each juvenile (independently) observes an environmental cue, as given in equation (3), and has its phenotype determined based on its genotype and the environmental cue; (*iv*) each juvenile has a probability *m* of migrating to a randomly chosen patch; and the cycle then returns to (*i*).

At the locus for *z* there are alleles *ζ_k_*, which we represent as real values limited to an interval. We are interested in situations where there is adaptively maintained genetic polymorphism at this locus. We think of the effects of the alleles as ‘genetic cues’, in the sense that they can provide statistical information to an individual about which habitat it is in (Leimar et al. 2006; Dall et al. 2015). In principle the alleles *ζ*_*k*_ can mutate, be selected, and evolve, but in order to aid the interpretation of our results, we make the simplification that there are two ‘fixed’ alleles, *ζ*_1_ and *ζ*_2_, that provide the genetic cues. The locus for the weight *α* in equation (1) can be seen as a ‘modifier’ locus, with alleles *α_k_*, that influence gene expression at the cue locus (note also that evolutionary changes of a modifier that is fully linked to *z* is equivalent to evolutionary changes of the genes at the locus for *z*). We represent the alleles *α*_*k*_ as real values in an interval. The phenotype in equation (1) is also influenced by the juvenile cue, mediated by the locus for the weight *β*, with alleles *β_k_*. In terms of plasticity, *β* is the slope of a reaction norm, and the alleles at the locus can be regarded as plasticity genes. We assume the loci are positioned in the order *z*, *α*, *β* along a chromosome, with *ρ*_*zα*_ the recombination rate between the cue locus and the modifier locus *α*, and *ρ*_*αβ*_ the rate between the modifier locus and the plasticity locus *β*.

The alleles at a locus are additive, producing diploid values as the sum of maternal and paternal allelic values. For instance, at the cue locus we have *z* = *ζ*_mat_ + *ζ*_pat_. The value *z* is referred to as a genetic effect or ‘genetic cue’, which can be polymorphic across habitats. For the loci giving the weights in equation (1), we are interested in cases where the modifier and slope effects, *α* = *α*_mat_ + *α*_pat_ and *β* = *β*_mat_ + *β*_pat_, are nearly monomorphic in the metapopulation, but evolving over the longer term.

### Two traits

We extend the situation above to two traits, *u*_1_ and *u*_2_, determined as

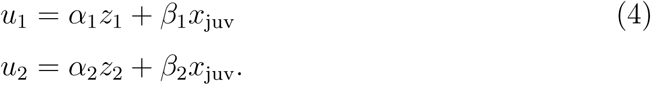

The genetic effects *z*_1_ and *z*_2_ are each determined by a locus with additive alleles, as in the case for a single trait above, and the juvenile environmental cue is given by equation (3). The modifiers *α*_1_, *α*_2_ and slopes *β*_1_, *β*_2_ are determined genetically by separate loci. The juvenile-to-adult survival in habitat *i* is given by

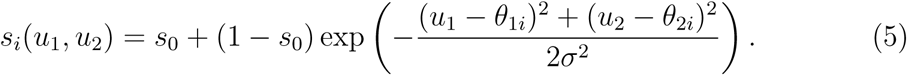

The loci are positioned in the order *z*_1_, *z*_2_, *α*_1_, *α*_2_, *β*_1_, *β*_1_ along a chromosome. Concerning recombination rates, *ρ*_*zz*_ is the recombination rate between the loci for the genetic effects *z*_1_ and *z*_2_, *ρ*_*zα*_ is the recombination rate between the locus for *z*_2_ and the locus for *α*_1_, and *ρ*_*αβ*_ is the recombination rate between neighboring loci for *α*_1_, *α*_2_, *β*_1_ and *β*_2_.

### Simulation model

For our individual-based simulations in Figs. 2 and 5, we started with a dimorphism at the locus for *z*, and allowed this dimorphism be maintained while *α* and *β* evolved. For some parameter values, for instance when *α* became close to 0, the dimorphism at the locus for *z* was not maintained. As mentioned, we used intervals for the allowed range of the values of alleles. For the simulations in Figs. 2 and 3 we used *ζ*_1_ = *−*0.4, *ζ*_2_ = 0.4 and the range [0.0, 4.0] for alleles at the loci for *α* and *β*. Mutational increments had a Laplace (reflected exponential) distribution with a standard deviation of 0.04, but allelic values were constrained to stay within the interval. The simulations were run for 100 000 generations with a mutation rate of 0.0050, to generate enough genetic variation for adaptation to proceed, followed by 100 000 generations with a mutation rate of 0.0001, to remove excess genetic variation. The simulations in Figs. 5 and 6 were performed in a similar way. The C++ source code for the computer programs used in this study is available from the Dryad repository (doi:xyz).

**Figure 2:**
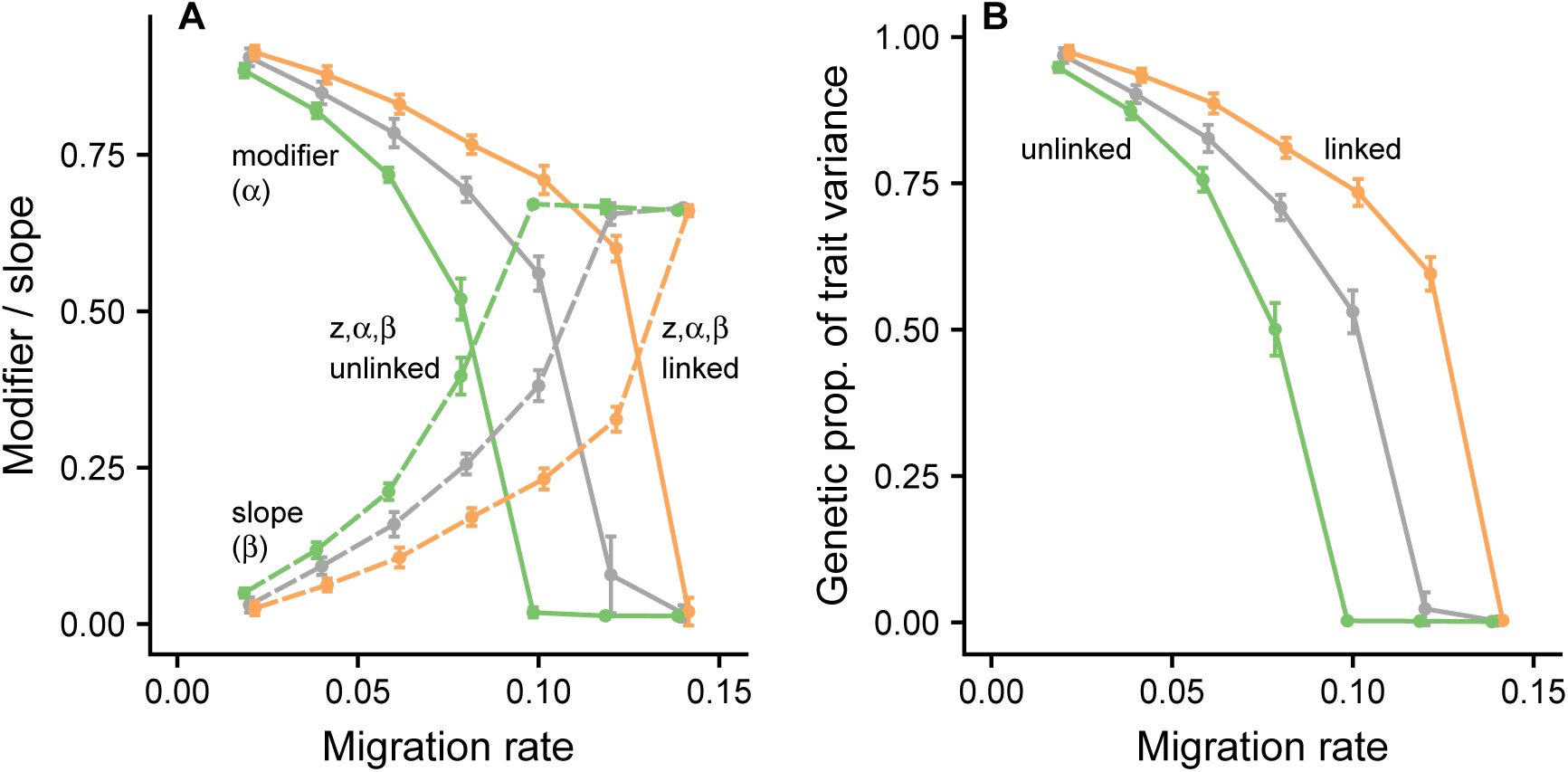
Phenotype determination for linked and unlinked genetic architectures, as a function of the rate of migration. Panel *A* shows how the epistatic modifier *α* (solid lines) of the genetic effect *z* and the slope *β* (dashed lines) of the reaction norm for the environmental cue *x*_juv_ depend on the migration rate *m* and on the genetic architecture. The mean SD over 10 replicate individual-based simulations is displayed. The left-hand (green) lines correspond to the case where the loci for *z*, *α* and *β* are all unlinked and the right-hand (orange) lines to the case where the three loci are tightly linked into a supergene. The lines between these (gray) correspond to an intermediate case where the loci for *z* and *α* are linked but the locus for *β* is unlinked to these. Panel *B* shows the genetic proportion of the partitioning of the variance of the phenotype *u* into genetic and plastic components. The genetic proportion is defined as the variance of the genetic component plus the covariance of the genetic and plastic components, divided by the total variance of the phenotype. Survival selection between habitats is given by equation (2) and the phenotype is determined as in equation (1). For the linked case, recombination rates are *ρ*_*zα*_ = *ρ*_*αβ*_ = 0.001, for the unlinked case *ρ*_*zα*_ = *ρ*_*αβ*_ = 0.5, and for the intermediate case *ρ*_*zα*_ = 0.001, *ρ*_*αβ*_ = 0.5. Other parameter values: *N*_*p*_ = 200, *K* = 100, *s*_0_ = 0.1, *σ* = 1.0, *θ*_1_ = *−*0.75, *θ*_2_ = 0.75, *ζ*_1_ = *−*0.4, *ζ*_2_ = 0.4, *σ*_juv_ = 0.5.

**Figure 3:**
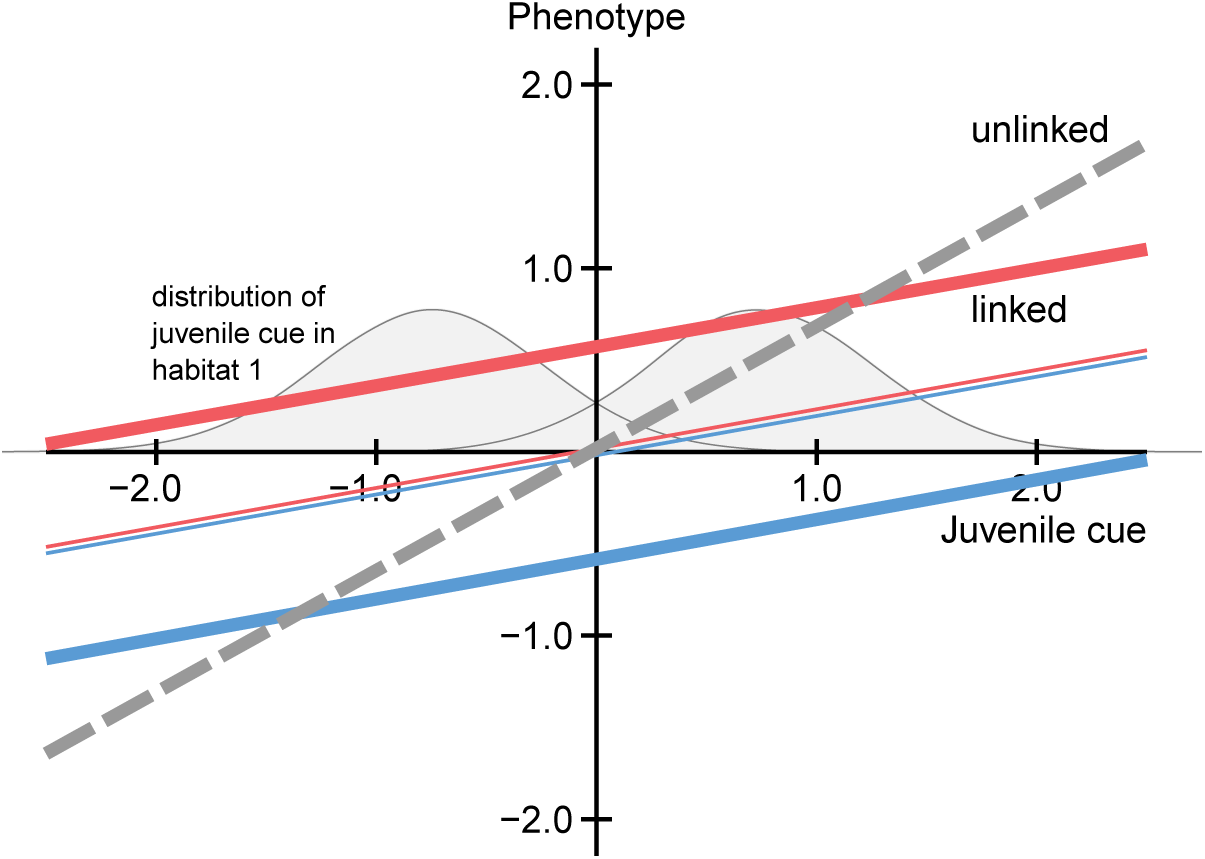
Example of the effect of genetic architecture (linked or unlinked) on phenotype determination. Mean reaction norms (with slope *β*) for habitat 1 specialists: thick and thin blue lines (slightly shifted up and down for clarity) represent individuals in habitat 1 with genotype *ζ*_1_*ζ*_1_ and *ζ*_1_*ζ*_2_ (with frequencies before migration of 0.76 and 0.22; line widths proportional to frequencies); and habitat 2 specialists: thick and thin red lines represent individuals in habitat 2 with genotype *ζ*_2_*ζ*_2_ and *ζ*_1_*ζ*_2_ (with frequencies 0.77 and 0.21); and for phenotypically plastic generalists: gray dashed line, slope and intercepts averaged over both habitats). For the generalist, the reaction norm is very similar between habitats (not shown), because *α* is small and *β* does not vary much, but the alleles *ζ*_1_ and *ζ*_2_ still segregate at the locus for *z*. The distributions of the juvenile environmental cue *x*_juv_ are shown lightly shaded for adult individuals in habitat 1 (left) and habitat 2 (right). The figure corresponds to the cases in fig. 2 for migration rate *m* = 0.10, with tightly linked loci for specialism and unlinked loci for plasticity.

**Figure 4:**
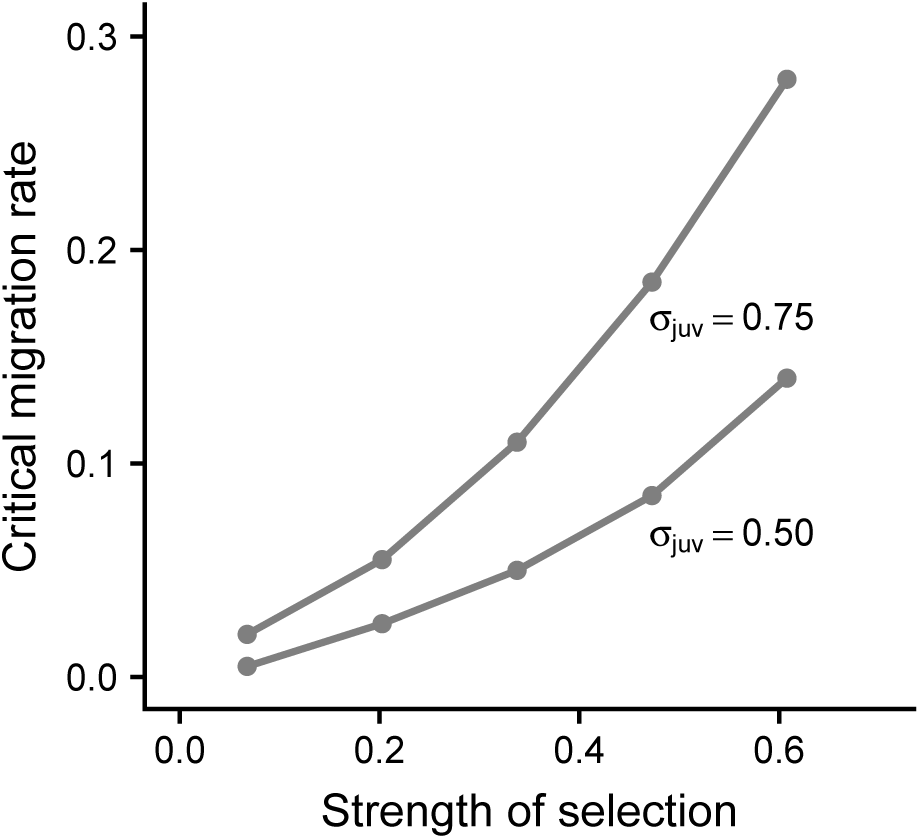
Critical migration rate, above which a genetic polymorphism in *z* is not selectively maintained, resulting in pure phenotypic plasticity. There is a single trait *u* and the loci for *z*, *α* and *β* are tightly linked. The critical rate is defined as the value of *m* for which the genetic proportion of the variance in *u* (see fig. 2B) is less than 0.01. The critical migration rate is shown as a function of the strength of selection in one habitat against a phenotype locally adapted to the other habitat, defined as 1 *s*_1_(*θ*_2_) = 1 *s*_2_(*θ*_1_) (see equation 2 for definition of *s*_*i*_). The points correspond to *s*_0_ = 0.9, 0.7, 0.5, 0.3, 0.1, and the lines are labeled with the juvenile environmental cue error, *σ*_juv_. The rightmost point on the line for *σ*_juv_ = 0.50 corresponds to the rightmost point for the linked case in fig. 2A, B. Other parameter values: *ρ*_*zα*_ = *ρ*_*αβ*_ = 0.001, *N*_*p*_ = 200, *K* = 100, *σ* = 1.0, *θ*_1_ = *−*0.75, *θ*_2_ = 0.75.

**Figure 5:**
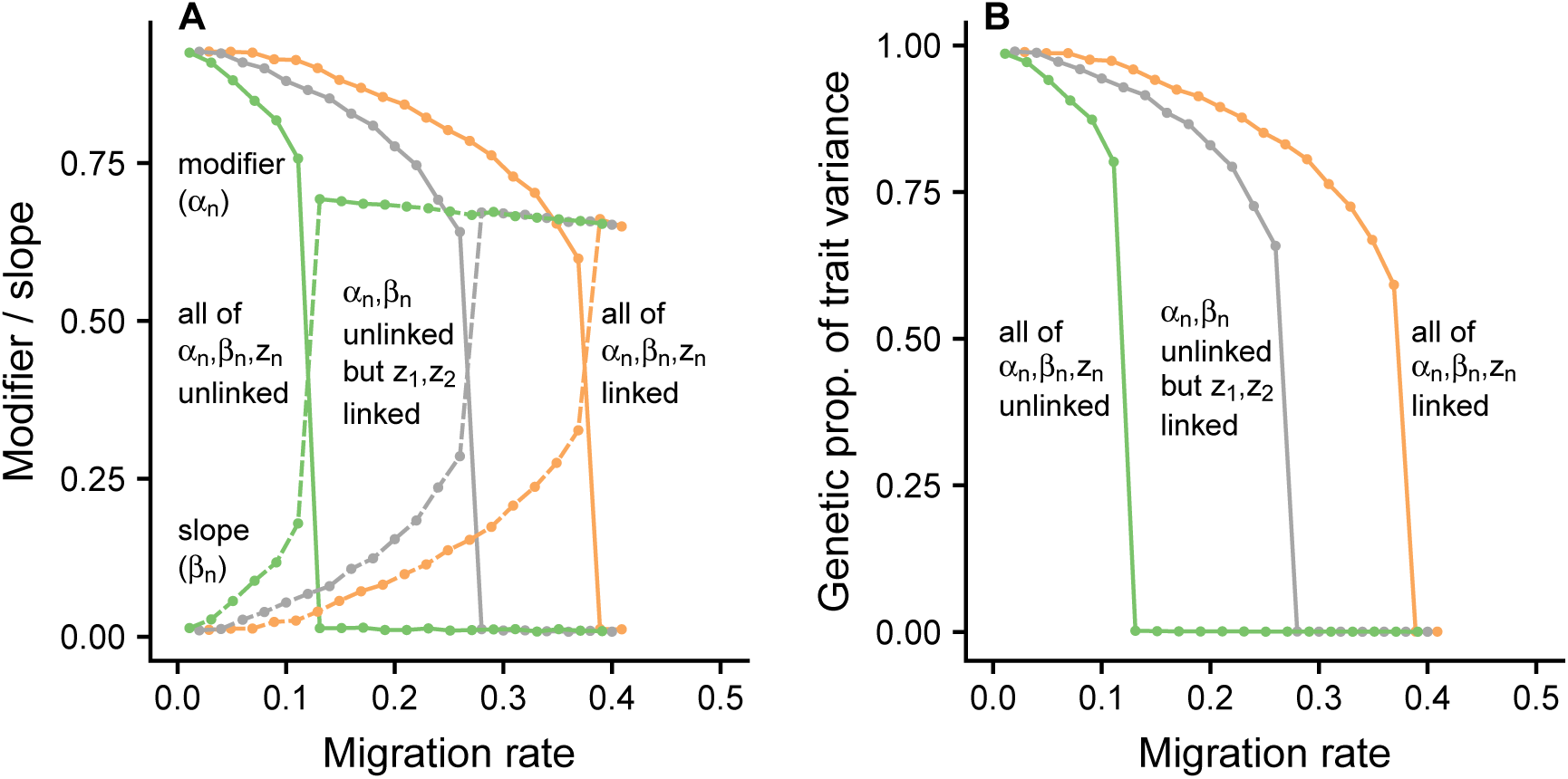
Phenotype determination for different genetic architectures, as a function of the rate of migration. Similar to fig. 2, but there are two traits, *u*_1_ and *u*_2_, each with optima that differ between the habitats. There are two genetic effect loci, one for each trait, and modifiers *α*_1_ and *α*_2_ for each of the genetic effects *z*_1_ and *z*_2_, as well as slopes *β*_1_ and *β*_2_ for the reaction norms of *u*_1_ and *u*_2_ for the juvenile cue *x*_juv_, following equation (4). Panel *A* shows how the mean modifier (*α*_1_ + *α*_2_)*/*2 and mean slope (*β*_1_ + *β*_2_)*/*2 depend on the migration rate *m* and on the genetic architecture. The solid lines show the mean modifier over 10 replicate of individual-based simulations, with the left-hand (green) line giving a case where the loci for the two genetic effects and the modifiers *α*_1_, *α*_2_, *β*_1_, *β*_2_ are all unlinked. The right-hand (orange) line shows the same thing, except that the six loci are tightly linked into a supergene. For the middle (gray) line, the two genetic effect loci are tightly linked, but the modifier and plasticity loci are unlinked from these and from each other. The dashed lines show the corresponding reaction norm slopes. The situation is symmetric between the traits, and the results for each trait separately are very similar to those shown here. Panel *B* shows the mean genetic proportion in the partitioning of the variance of the phenotypes *u*_1_ and *u*_2_ into genetic and plastic components. Survival selection between habitats is given by equation (5). For the linked case, recombination rates are *ρ*_*zz*_ = *ρ*_*zα*_ = *ρ*_*αβ*_ = 0.001, and for the unlinked case *ρ*_*zz*_ = *ρ*_*zα*_ = *ρ*_*αβ*_ = 0.5. Other parameter values: *N*_*p*_ = 200, *K* = 100, *s*_0_ = 0.1, *σ* = 1.0, *θ*_11_ = *θ*_21_ = *−*0.75, *θ*_12_ = *θ*_22_ = 0.75, *σ*_juv_ = 0.5.

**Figure 6:**
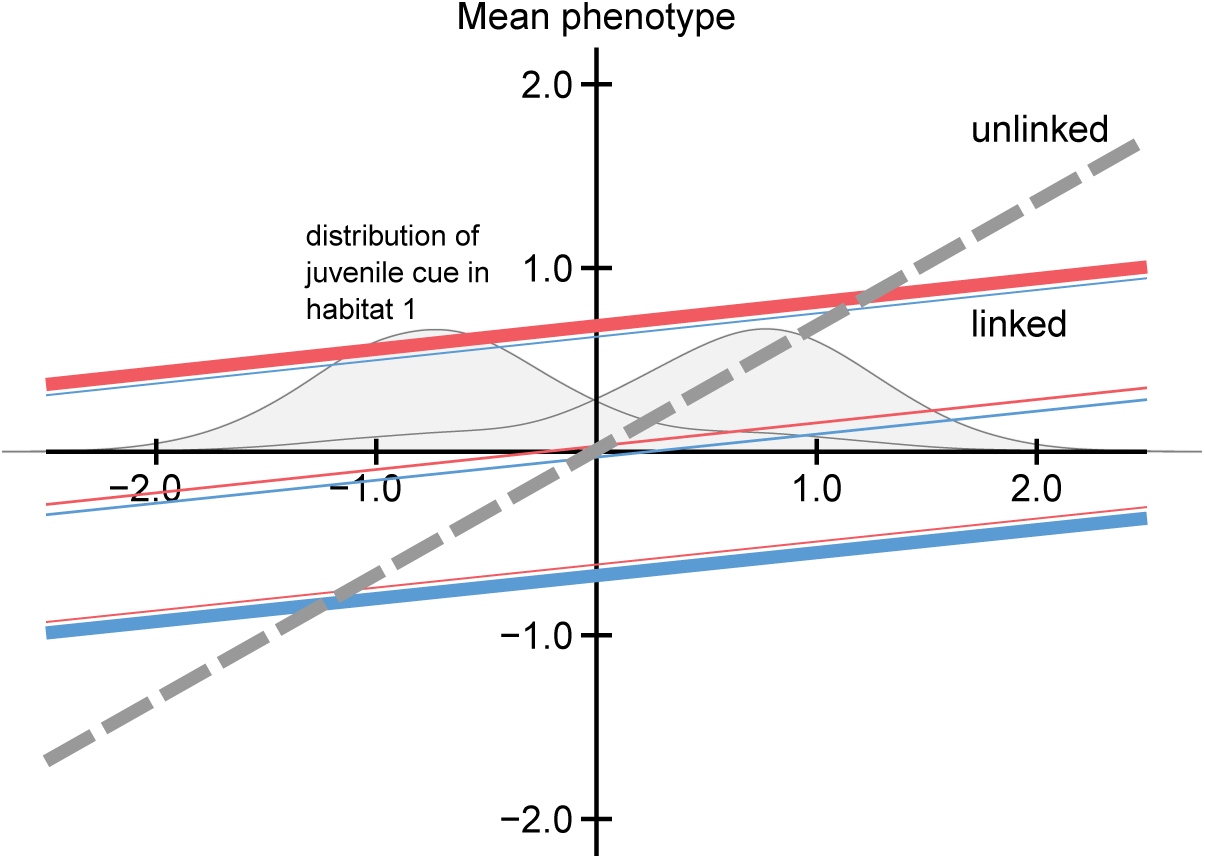
Example of the effect of genetic architecture (linked or unlinked) on pheotype determination. Mean reaction norms (with slope (*β*_1_ + *β*_2_)*/*2) for habitat 1 specialists: thick and thin blue lines (slightly shifted up and down for clarity) represent individuals in habitat 1 with genotype *ζ*_1_*ζ*_1_, *ζ*_1_*ζ*_2_ and *ζ*_2_*ζ*_2_ at each of the two genetic effect loci (with frequencies after migration of 0.74, 0.15 and 0.10; line widths proportional to frequencies); and habitat 2 specialists: thick and thin red lines represent individuals in habitat 2 with genotype *ζ*_2_*ζ*_2_, *ζ*_1_*ζ*_2_ and *ζ*_1_*ζ*_1_ at each of the two genetic effect loci (with frequencies 0.73, 0.16 and 0.11); and for phenotypically plastic generalists: gray dashed line, slope and intercepts averaged over both habitats). For the liked case (specialist), the genotypes at the loci for *z*_1_ and *z*_2_ are highly correlated, both among habitats (correlation of genetic effects: 0.999) and within habitats (0.998). For the generalist, the reaction norm is very similar between habitats (not shown), because the *α*_*n*_ are small and the *β*_*n*_ do not vary much, but the alleles *ζ*_1_ and *ζ*_2_ still segregate at the locus for *z*_2_, whereas in this example *z*_1_ is fixed for *ζ*_2_. The distributions of the juvenile environmental cue *x*_juv_ are shown lightly shaded for adult individuals in habitat 1 (left) and habitat 2 (right). The figure corresponds to the cases in fig. 5 for migration rate *m* = 0.24, with tightly linked loci for specialism and unlinked loci for plasticity.

## Results

The effect of genetic architecture on local adaptation and phenotypic plasticity is illustrated in Figs. 2 and 3, with data from individual-based simulations. There is a single trait *u*, with optimal survival at trait value *θ*_1_ and *θ*_2_ in habitat 1 and 2 (equation 2). The determination of the phenotype is given by *u* = *αz* + *βx*_juv_, where *z* is a genetic effect, *α* is an epistatic modifier of *z*, *x*_juv_ is an environmental cue (equation 3), and *β* is a plasticity effect, giving the slope of a reaction norm (equation 1). Each of *z*, *α*, and *β* is determined by a single diploid locus with additive allelic effects, and we are comparing the case where the loci are tightly linked into a supergene with that where they are all unlinked (Figs. 2, 3). As seen in fig. 2, for intermediate rates of migration between habitats the genetic architecture strongly influences the contributions of genetic polymorphism and plasticity to variation in *u*. For tightly linked loci, the genetic contribution to the variation is larger than for unlinked loci, and the reverse is true for the contribution from plasticity.

The ecological genetic conflict is further exemplified by the reaction norms for migration rate *m* = 0.10 between local populations (corresponding to a migration rate of 0.05 between habitats), which are shown in fig. 3, together with the distributions of the environmental cue that adults in the different habitats observed as juveniles. For the linked case, there are reaction norms with shallower slopes, with different mean intercepts for individuals in habitats 1 and 2 with different genotypes (red and blue lines in fig. 3 represent habitat 1 and 2 specialists, cf. fig. 1C). There is genetic variation in *z* in each habitat: there are two alleles, each better adapted to one of the habitats, giving rise to alternative homozygotes and heterozygotes, with different frequencies in the habitats (in principle, these genes can evolve, and a balance between mutation, selection and drift can maintain variation around each of them). For the unlinked case, there is single reaction norm with steeper slope (dashed line in fig. 3), corresponding to a phenotypically plastic generalist. Note that the only difference in model parameters between the linked and unlinked cases is the genetic architecture, demonstrating that ecological genetic conflict can have a pronounced influence on phenotype determination.

The issue of divergence of evolutionary interests between specialism and plasticity hinges on whether genes tightly linked to one of the alleles at the polymorphic locus for *z*, adapted to one of the habitats, has an appreciable chance of recombining to become associated with an allele locally adapted to the other habitat, as well as migrating to that habitat. The way this can happen is if a modifier allele occurs in a heterozygote between alleles at the locus for *z*, each adapted to different habitats. The strength of selection against such a heterozygote influences the chance for the modifier allele to recombine to the other locally adapted allele. For the linked case shown in fig. 3, this chance is small, illustrating that genes for specialism have their evolutionary future mainly in their own habitat. While studying between-habitat genetic polymorphism, Bengtsson (1985) and Barton and Bengtsson (1986) introduced the concept of an effective migration rate for a neutral locus that is linked to a selected, genetically polymorphic locus. For instance, using equation (4) in Yeaman and Whitlock (2011), and ignoring the effects of plasticity, we find an effective migration rate of 0.0002 for a linkage of *ρ* = 0.001 to *z* (fig. 3), so for such genes for specialism the two habitats are fairly isolated from each other.

An alternative and more informative way of showing how the evolutionary interest varies with the degree of linkage to a between-habitat polymorphism is to examine how the reproductive value for a modifier of being associated (linked) with an allele adapted to one or the other habitat depends on the rate of recombination. We have performed a numerical analysis of a model with a very large population in each habitat (see supporting information for model description), but otherwise similar to the simulation model with results in Figs. 2 and 3. The results of the numerical analysis, which takes into account plasticity, are given in Table A1 and fig. A1. The outcome of then analysis using reproductive values is less extreme but qualitatively similar to the consideration of effective migration rates. As seen in Table A1, for *m* around 0.1 (*m*_12_ around 0.05) and with modifiers tightly linked to *z*, the reproductive value of being associated with the locally adapted allele at the genetic effect locus is around four times higher than that of being associated with the other allele, whereas these values are nearly equal for loosely linked modifiers.

In any case, for migration rate above a critical value, phenotype determination for the linked case (as well as for the unlinked case) is dominated by plasticity, because the modifier *α* in equation (1) approaches zero. For instance, in fig. 2 the critical migration rate is *m* = 0.14. The critical migration rate for a wider range of parameters is shown in fig. 4. In general, stronger selection between habitats and less accurate juvenile environmental cues favor genetic polymorphism in *αz*, and thus a higher value of the critical migration rate (fig. 4).

The emergence of genomic islands of divergence has been modeled as several linked genes of smaller effect that add up to a bigger effect for a particular trait (e.g., Yeaman and Whitlock 2011; Yeaman 2013). In order to make contact with this work, we performed individual-based simulations with 100 linked loci, with the alleles at each locus constrained to have small effects, and with parameters similar to those in fig. 2. Plasticity was prevented from evolving in this simulation. As shown in fig. A2, based on the between-habitat *F*_ST_ for each locus, an island of divergence spanning around 15 loci emerged.

For two traits, *u*_1_ and *u*_2_, each with different optima in the habitats, as given by equation (5), we again find a pronounced influence of genetic architecture on the relative importance of genetic polymorphism and plasticity (fig. 5, 6). For each trait, *u*_1_ and *u*_2_, there is a separate genetic effect, *z*_1_ and *z*_2_, coded by one locus, with modifier *α*_1_ and *α*_2_ and reaction norm slope *β*_1_ and *β*_2_, but the same juvenile environmental cue *x*_juv_ for both reaction norms, as given in equation (4). Three cases are illustrated in fig. 5, one where all loci are linked, another where the two genetic effect loci are linked and the loci for *α*_1_, *α*_2_, *β*_1_ and *β*_2_ are unlinked from each other and from the genetic effect loci, and a third case where all loci are unlinked. From this figure, and the example in fig. 6, it appears that the influence of genetic architecture is qualitatively similar but even stronger for a two-trait syndrome compared to a single trait. Again, we find that for each trait several genes of smaller effect can add up to a bigger effect, as shown in fig. A3.

For the two-trait syndrome, we explored the evolution of linkage using individual-based simulations. Instead of specifying the recombination rates *ρ*_*zz*_, *ρ*_*zα*_ and *ρ_αβ_*, we let these be coded by three loci. We found that tight linkage between the two polymorphic effect loci *z*_1_ and *z*_2_ promptly evolved (i.e., *ρ*_*zz*_ became close to zero; Table 1), so these loci emerge as an island of divergence.

**Table 1:** Evolution of linkage for two-trait simulations similar to fig. 5. There are 9 loci along a chromosome, coding for *z*_1_, *z*_2_, *ρ*_*zz*_, *ρ*_*zα*_, *ρ_αβ_*, *α*_1_, *α*_2_, *β*_1_, and *β*_2_, and the table gives averages (±SD for recombination rates) in the population after 200000 generations. The loci for the recombination rates are tightly linked to the locus for *z*_2_, in order to maximize the chances of the evolution of tighter linkage to the polymorphic complex *z*_1_ - *z*_2_. The recombination rate *ρ*_*zz*_ between *z*_1_ and *z*_2_ evolved towards tight linkage, but the other recombination rates reached intermediate average values, with broad distributions, as illustrated in fig. 7.

| $m$ | $\rho_{zz}$ | $\rho_{z\alpha}$ | $\rho_{\alpha\beta}$ | $\alpha_1$ | $\alpha_2$ | $\beta_1$ | $\beta_2$ |
| --- | --- | --- | --- | --- | --- | --- | --- |
| 0.12 | $0.0029 \pm 0.0071$ | $0.205 \pm 0.300$ | $0.209 \pm 0.110$ | 0.891 | 0.897 | 0.073 | 0.063 |
| 0.18 | $0.0019 \pm 0.0041$ | $0.028 \pm 0.022$ | $0.243 \pm 0.115$ | 0.871 | 0.856 | 0.098 | 0.113 |
| 0.24 | $0.0008 \pm 0.0060$ | $0.251 \pm 0.200$ | $0.335 \pm 0.060$ | 0.688 | 0.673 | 0.238 | 0.244 |

However, for *α*_1_, *α*_2_, *β*_1_ and *β*_2_ we did not find notable selection for either tighter or looser linkage to the *z*_1_ - *z*_2_ complex. Considerable genetic variation for the recombination rates *ρ*_*zα*_ and *ρ*_*αβ*_ persisted in the population, perhaps as a result of mutation-drift balance (see Table 1 and fig. 7 for illustration of these simulations). Overall, the outcome for the modifiers *α*_1_, *α*_2_ and plasticity slopes *β*_1_, *β*_2_ was similar to the middle (gray) case in fig. 5, with tightly linked *z*_1_ and *z*_2_ and unlinked loci for modifiers and slopes.

**Figure 7:**
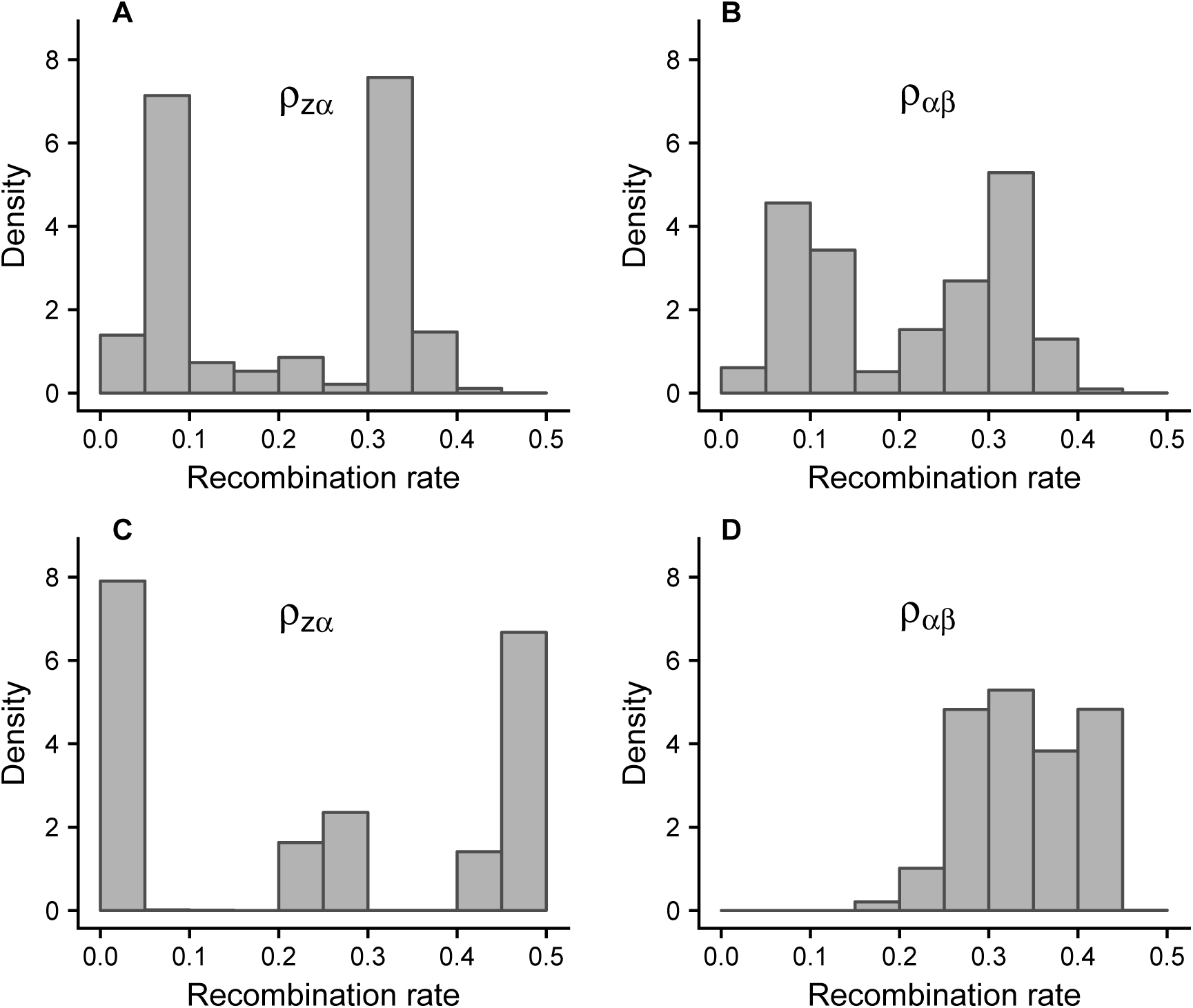
Distribution of recombination rates, over individuals in the population, from the simulations for *m* = 0.12 and *m* = 0.24 reported in Table 1. Panels *A* and *B* show *ρ*_*zα*_ and *ρ*_*αβ*_ for the case with *m* = 0.12 in Table 1, and *C* and *D* show the same for the case with *m* = 0.24. Overall, there seems not to be a tendency for evolution of either very low or very high recombination rates.

## Discussion

Both habitat specialism and plasticity are well-studied phenomena (van Tienderen 1991, 1997; West-Eberhard 2003; DeWitt and Langerhans 2004; Richards et al. 2006; Griffith and Sultan 2012), but the perspective of divergence of evolutionary interests has traditionally not been applied. By examining ecological genetic conflict, we have identified phenomena that were not studied before. Compared to previous models of the evolution of genomic islands of divergence, the major new aspect of our work is that we study phenotypic plasticity together with genetic polymorphism, and that we interpret our results in terms of genetic conflict, or divergence of evolutionary interests, between genes for specialism and phenotypically plastic generalism. We find that the rate of recombination between genetic effect, modifier and plasticity loci influences the evolutionary outcome, with more plasticity and less genetic polymorphism for unlinked loci, in particular for intermediate migration rates (Figs. 2 and 5).

Our explanation is that modifier and plasticity genes unlinked to a polymorphic genetic effect locus favor phenotypes that are less specialized to a particular habitat compared to tightly linked genes, because unlinked genes become adapted to exist in all habitats. Tightly linked modifier and plasticity genes, on the other hand, are selected to perform well mainly in one of the habitats, even at the expense of performance in another habitat. Thus, a modifier or plasticity allele tightly linked to an allele at a polymorphic locus can become concentrated to one of the habitats, with the other habitat acting as a sink, to which little adaptation takes place (Holt and Gaines 1992; Kawecki 1995). One way of quantifying this effect is as a low effective migration rate for loci tightly linked to a genetic polymorphism (Barton and Bengtsson 1986; Yeaman and Whitlock 2011; Aeschbacher et al. 2017), and another and perhaps more informative approach is to compute reproductive values of modifiers, as we have done (Table A1). Note that an allele at a polymorphic genetic effect locus does have a future also when present as a heterozygote in the ‘wrong habitat’, because migration can transport it back to the other habitat. Thus, migration makes the distinction between linked and unlinked genetic architectures a matter of degree rather than kind.

In fact, the general pattern of variation of the modifier *α* and plasticity slope *β* with the migration rate *m* is qualitatively similar for different genetic architectures, with a shift from mainly genetic polymorphism to mainly phenotypic plasticity as *m* increases (Figs. 2, 5, S1). One way of explaining this shift is in terms of the statistical information about the habitat that is contained in the ‘genetic cue’ *z* in comparison with the environmental cue *x*_juv_ (Leimar et al. 2006; Leimar and McNamara 2015; Dall et al. 2015). Tufto (2000) provides a discussion of earlier papers dealing with this topic. For higher values of *m*, gene frequency differences between habitats are smaller, thus being less statistically informative about the habitat compared to the environmental cue *x*_juv_. An optimal phenotype determination strategy will therefore put less emphasis on the genetic and more on the environmental cue for higher values of *m*. For high enough rates of migration, and provided that environmental cues are sufficiently accurate, phenotypic plasticity dominates completely, as illustrated by simulations in fig. 4. For a much simpler model with binary cues, inspired by the work of Sultan and Spencer (2002), an analytical solution is possible, leading to qualitative similar results (see equation 4 and fig. 5 in Leimar et al. 2006). Note also that, if migration rates are not too high, a generalist strategy of phenotype determination can make use of the information form a polymorphic genetic cue, provided that the polymorphism is selectively maintained. The unlinked case with *m* = 0.06 in fig. 2 and that with *m* = 0.10 in fig. 5 are examples of this outcome.

Our conclusion that the evolution of genomic islands of divergence is favored by a combination of migration and divergent selection between habitats is in qualitative agreement with previous theoretical analyses (e.g., Aeschbacher et al. 2017), including our result (fig. 4) that there is a critical migration rate above which migration dominates over selection (e.g., Yeaman and Whitlock 2011). Note, however, that our analysis examines how genetic polymorphism balances with phenotypic plasticity, rather than with genetic drift. The general idea that migration and divergent selection promote genomic islands of divergence also has empirical support (Samuk et al. 2017).

Our main result, that reaction norm slopes can depend on the genetic architecture (Figs. 2, 3, 5, 6, S1), is new and there appear to be no empirical data directly examining this question. It is known that ecotypic traits differ in how they are determined, with the variation in some traits being mainly genetic and in other traits mainly plastic (Lucek et al. 2014), but the possible influence of genetic architecture is unknown. There are observations showing that plasticity can decrease during the formation of an ecotype (Hasan et al. 2017), but the genomic basis of the reduction in plasticity is not known. Also, a study of so called expression quantitative trait loci (eQTLs) shows that ‘distant’, *trans*-regulatory changes on average had different effects than ‘local’, *cis*-regulatory changes, and were also more responsive to the environment (Ishikawa et al. 2017), which is at least suggestive of an influence of genetic architecture on trait expression.

In our investigation of the evolution of recombination, for a two-trait situation, we found that low recombination between the polymorphic loci for *z*_1_ and *z*_2_ readily evolved (Table 1), corresponding to an island of divergence, and this is in accordance with the traditional understanding of such situations (Pinho and Hey 2010; Via 2012). On the other hand, we did not detect selection for either tighter or looser linkage between the polymorphic loci and epistatic modifiers or plasticity loci (Table 1). The question appears not to have been analyzed previously, but perhaps other factors that could influence genetic architecture, such as inversions or a tendency towards *cis*-regulatory influences, play a greater role in determining the linkage.

The idea that genes occurring in linked clusters, whether in ecotypes or in other contexts, share an evolutionary interest by being transmitted together, points to the possibility that genetic conflict is of importance for many instances of supergenes and co-adapted gene complexes (Schwander et al. 2014; Thompson and Jiggins 2014; Charlesworth 2016). The ‘genomic islands’ found in microorganisms (Hacker and Carniel 2001; Dobrindt et al. 2004) might have a similar explanation. In conclusion, by applying traditional ideas of genetic conflict to genomic islands of divergence in ecotypes, we have extended the concept of genetic conflict to an ecological context and produced new and fundamental results about the balance between genetic local adaptation and phenotypic plasticity. We hope that our work can inspire further empirical investigation of the genomics of phenotypic plasticity of ecotypes.

## Appendix A

### Numerical analysis

Our approach here shows similarity to the numerical analysis by Leimar *et al.* (2016). The main aim of this analysis is illustrate the divergence of evolutionary interests between tightly linked and unlinked modifiers of a polymorphic genetic effect locus, through the use of reproductive values, as illustrated in Table A1. We also show how the modifier *α* and the slope *β* vary as the rate of recombination between these loci and the genetic effect increases from 0 to 0.5 (fig. A1).

Let habitat *i*, *i* = 1, 2, support a large population of size *n*_*i*_ and let *m*_*ij*_ be a rate of migration to habitat *i* from habitat *j*, in the sense that, after migration, the respective proportions *m*_11_ and *m*_12_ of individuals in habitats 1 originate from habitat 1 and 2, and similarly in habitat 2. We are mostly interested in the symmetric case where *n*_1_ = *n*_2_, *m*_11_ = *m*_22_ and *m*_12_ = *m*_21_ The life cycle of individuals is a version of that in the main text: (*i*) within-habitat random mating, forming *n*_*i*_ offspring in habitat *i*, conceptualized as random unions from a pool of gametes, drawn from the adults in the habitat (after which the adults die); (*ii*) each juvenile (independently) observes an environmental cue, as given in equation (3), and has its phenotype determined based on its genotype and the environmental cue; (*iii*) each juvenile has a probability *m*_*ij*_*n*_*i*_/*n*_*j*_ of migrating from its habitat *j* to habitat *i*; (*iv*) selection, with survival in habitat *i* as a function of phenotype *u* as in equation (2); and the cycle then returns to (*i*).

Let us use notation like *ζ*_*k*_ to denote alleles at the locus for *z*. We take (*i*) as our census point, and let *p*_*ik*_ be the frequency among the gametes (that form the next generation) of allele *ζ*_*k*_ in habitat *i*. If we order the gametes as maternal-paternal, the genotype frequencies among the offspring at the census point in habitat *i* are *p_ik_p_il_*. Concerning environmental cues, note that the mean cue in habitat *i* is *θ*_*i*_, according to equation (3). The survival in habitat *i* of individuals with genotypes with alleles *ζ*_*k*_ and *ζ*_*l*_ who have observed the juvenile cue in habitat *j* becomes

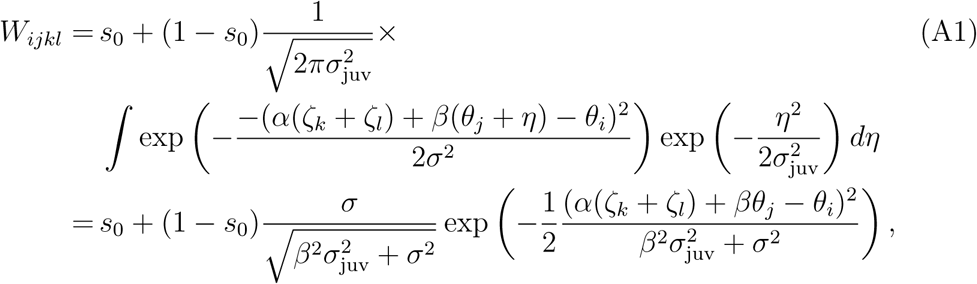

where the integration variable *η* represent the environmental cue error. Note that we have the symmetry *W*_*ijkl*_ = *W*_*ijlk*_. Define an average survival as

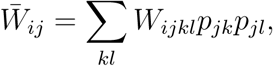

we get the genotype frequencies at the end of phase (*iv*) as

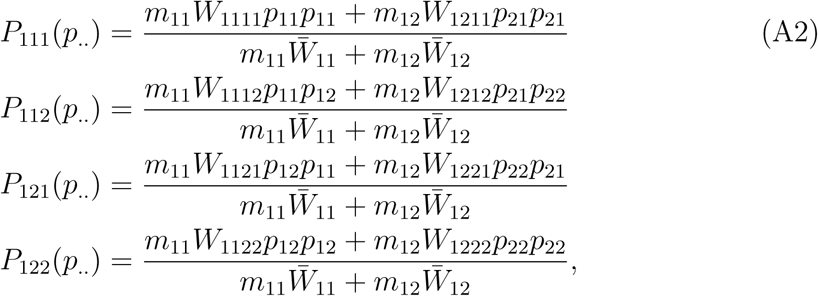

in habitat 1, and

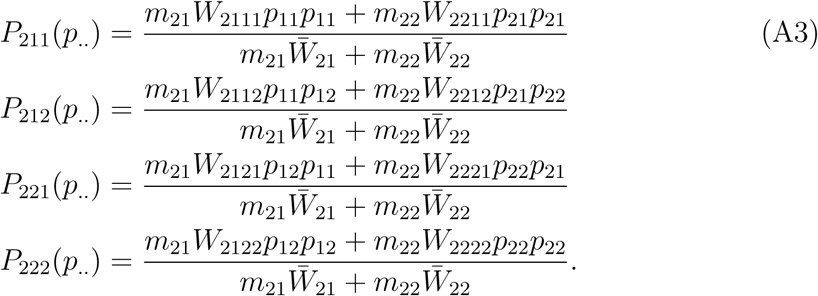

in habitat 2. The notation *P*_*ikl*_(*p_.._*) means that there is a dependence on the allele frequencies: *p*_.._ = (*p*_11_*, p*_21_*, p*_12_*, p*_22_). Again, we have the symmetry *P*_*ikl*_(*p_.._*) = *P*_*ilk*_(*p_.._*), and the index combination *kl* means that *k* is the maternal and *l* the paternal allele. From one generation to the next, we then have the following iteration for the allele frequencies at the census point:

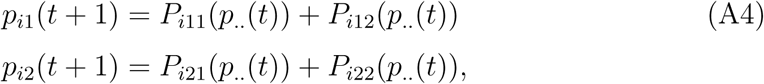

where we have taken into account the symmetry *P*_*i*12_ = *P*_*i*21_. We can note that *p*_*i*1_(*t* + 1) + *p*_*i*2_(*t* + 1) = 1, as it should, so we only need the equation for *p*_*i*1_. The iteration (A4) can be used to determine numerically the equilibrium allele frequencies for a given situation, as is done in Table A1. In the following, we let *p*_*ik*_ denote such an equilibrium.

### Mutant invasion

We now consider a rare mutant modifier, that modifies either *ζ*_1_, *ζ*_2_, *α* or *β*, and that has a rate of recombination *ρ* with the polymorphic locus for *z*. To make it simple, we assume that a modifier changes either *ζ*_1_ to 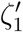, or *ζ*_2_ to 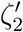, when linked to that allele, or modifies *α* to *α′* or *β* to *β′*. Let 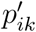 bet the frequency in habitat *i* of a mutant modifier linked to allele *k*, with 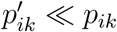, and let 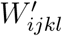 be the modified survival where the modifier is linked to allele *l*. Here, we do not distinguish maternal and paternal origin. Similar to equations (A2, A3), we have the first-order terms in mutant frequencies as

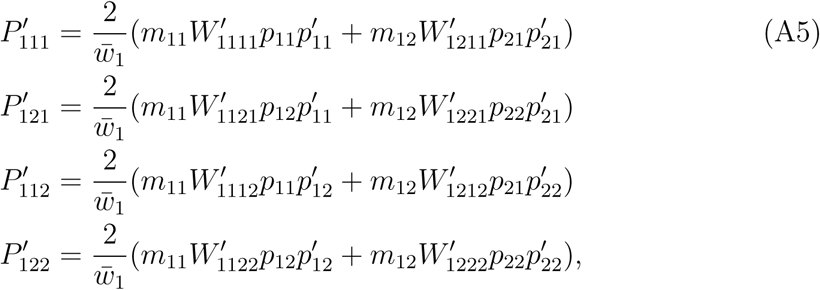

and

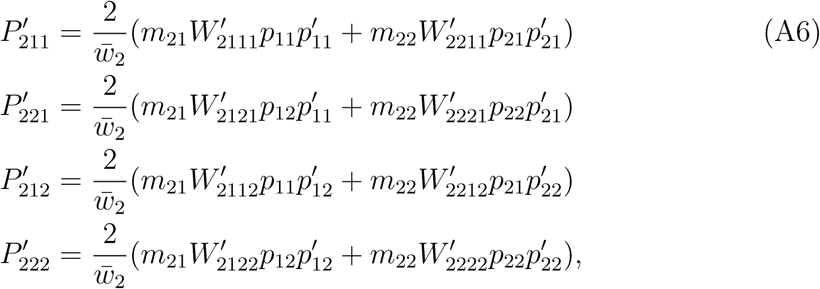

where we used the notation 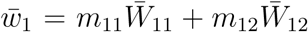 and 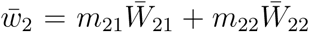. These represent mutant heterozygote genotypes surviving to the census point, ready to produce gametes for next generation: 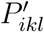 is the frequency of mutant heterozygotes in habitat *i* where the mutant modifier is linked to the *l* allele. Recombination gametes from 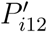 and 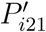 can transfer the mutant modifier to become linked to the other allele at the locus for *z*. Using this, the iteration from one generation to the next for the 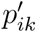 becomes:

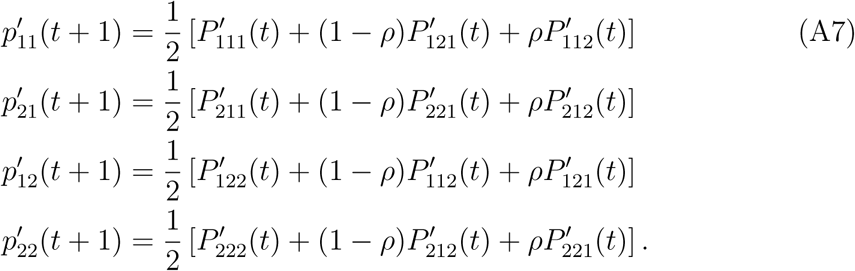

We can write the mutant population projection as

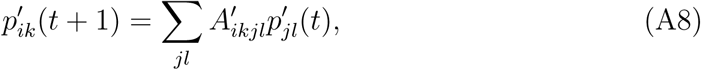

where 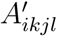 is the population projection matrix. We get

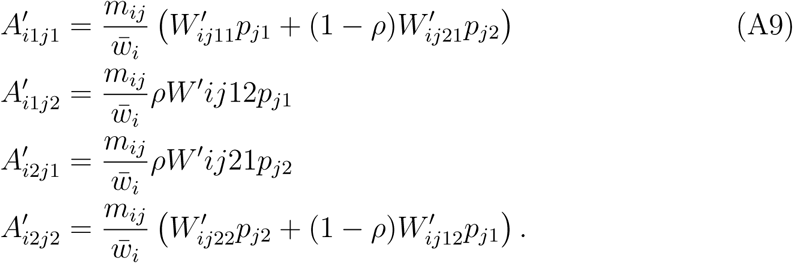

The mutant projection is a 4 *×* 4 matrix, and each line of equation (A9) represents a partitioning of this matrix into 2 *×* 2 sub-matrices.

### Invasion fitness

The leading eigenvalue *λ* of the matrix **A**′, with elements 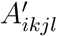, or rather its logarithm, log *λ*, gives the mutant invasion fitness. For the case where the mutant is equal to the resident, we have *λ* = 1, with (*p*_11_*, p*_21_*, p*_12_*, p*_22_) as right eigenvector and the reproductive values (*v*_11_*, v*_21_*, v*_12_*, v*_22_) as left eigenvector. Furthermore, the mutant can invade if *λ >* 1.

We developed a C++ program that follows a path of small steps through either *ζ*_1_*ζ*_2_–space, or *αβ*–space, each of which increases the invasion fitness, until reaching an accurate approximation of the equilibrium. We first put *α* = 1 and *β* = 0 and looked for an equilibrium dimorphism *ζ*_1_*ζ*_2_. We then retained this dimorphism and let *α* and *β* evolve to an equilibrium, for different values of the rate of recombination *ρ* between the locus for *ζ*_1_*ζ*_2_ and the loci for *α* and *β*. In this analysis, we made the assumption that *α* and *β* are tightly linked to each other. The result of the analysis is presented in Table A1. An important point of the analysis appears in the final column, giving the ratio *v*_11_*/v*_12_ of the reproductive value for a small-effect modifier (in the limit of being neutral) of being associated with the locally favoured allele *ζ*_1_ to being associated with the other allele *ζ*_2_. This ratio expresses how much a small increase in survival in one habitat is weighed against a corresponding decrease in survival in the other habitat.

**Table A1:**
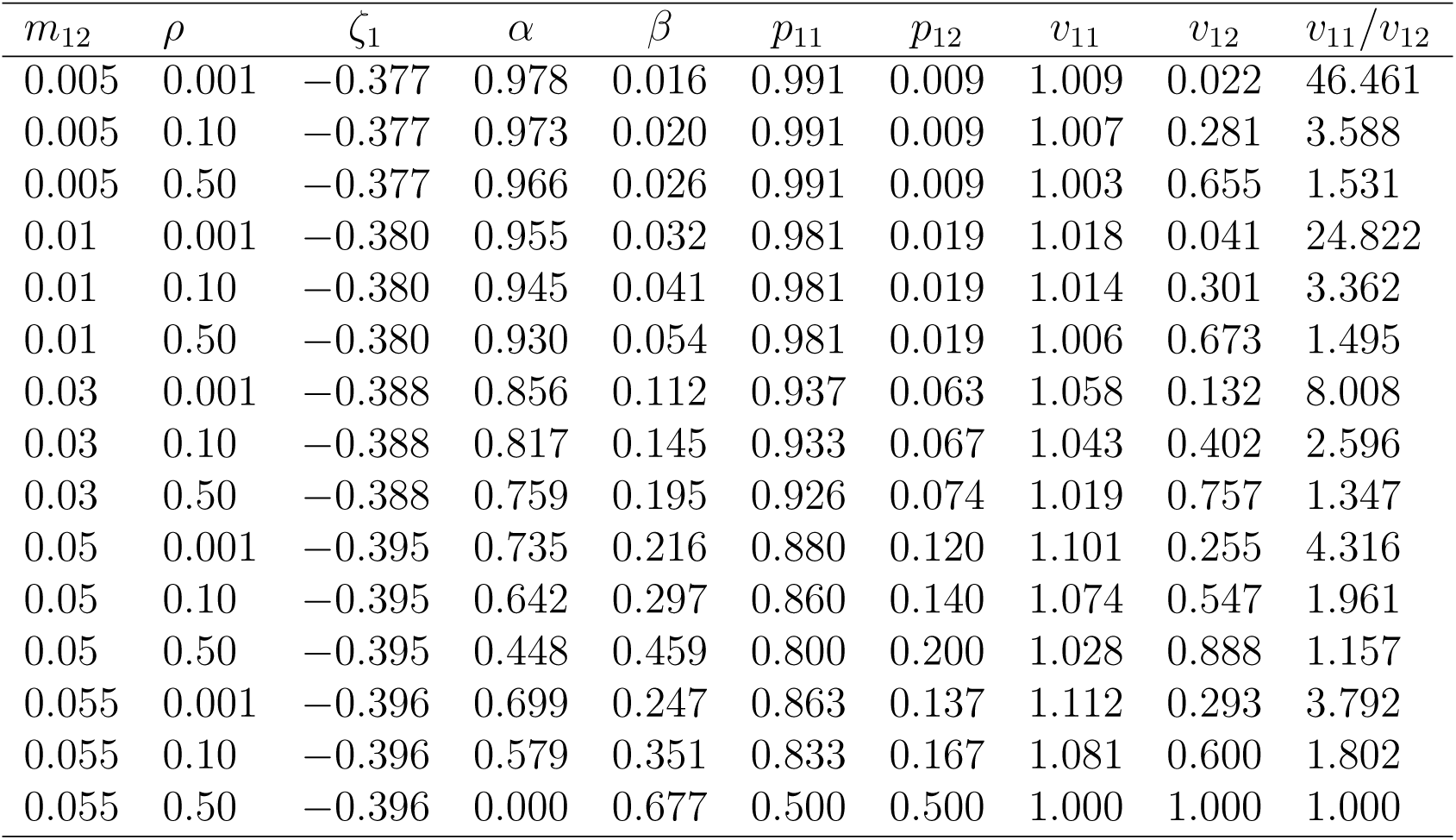
Numerical analysis of the alternative model. It is similar to the simulation model explored in the main text, with results in figs. 2 and 3. The main difference is that, in the alternative model, each habitat supports a single very large population, instead of several smaller local populations. Phenotype determination follows equation (1) with survival in each habitat given by equation (2) and environmental cues as in equation (3). The rate of migration between habitats is denoted *m*_12_ (with *m*_21_ = *m*_12_) and corresponds to *m/*2 in the model in the main text. The table shows the rate of between-habitat migration *m*_12_, the rate of recombination *ρ* between the genetic effect locus and the loci for *α* and *β*, the value *ζ*_1_ of the allele adapted to habitat 1 at the genetic effect locus (with *ζ*_2_ = *−ζ*_1_), the equilibrium values of the modifier *α* and the slope *β*, the frequencies *p*_11_ and *p*_12_ in habitat 1 of the alleles *ζ*_1_ and *ζ*_2_ at the time of reproduction, and the reproductive values *v*_11_ and *v*_12_ of small-effect mutant modifiers, with linkage *ρ* the the genetic effect locus. The value *v*_11_ applies when the mutant modifier is linked to the locally adapted allele *ζ*_1_ and *v*_12_ when linked to the alternative allele *ζ*_2_. The final column gives the ratio of the reproductive values, which indicates how strongly modifications that improve performance in habitat 1 are favored. Note that the situation is symmetric, with *p*_21_ = *p*_12_, *p*_22_ = *p*_11_, *v*_21_ = *v*_12_ and *v*_22_ = *v*_11_. Other parameter values: *s*_0_ = 0.1, *σ* = 1.0, *θ*_1_ = *−*0.75, *θ*_2_ = 0.75, *σ*_juv_ = 0.5.

**Figure A1:**
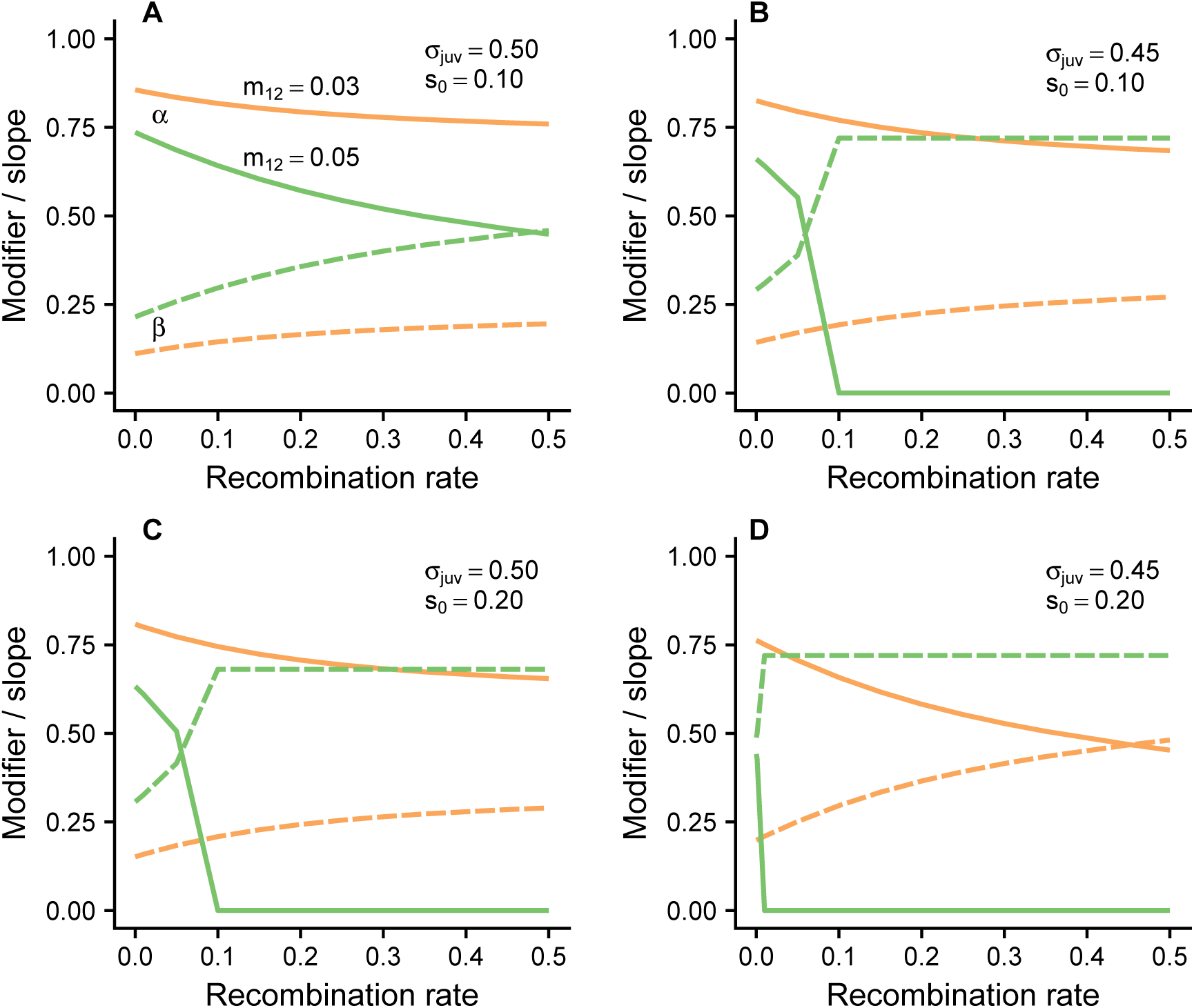
Numerical analysis of the alternative model. The panels show the modifier *α* (solid lines) and slope *β* (dashed lines) for different values of the parameters *m*_12_ = *m*_21_ (orange and green lines), *σ*_juv_, and *s*_0_, as a function of the recombination rate *ρ* between between the genetic effect locus and the loci for *α* and *β*. Other parameter values: *σ* = 1.0, *θ*_1_ = *−*0.75, *θ*_2_ = 0.75.

**Figure A2:**
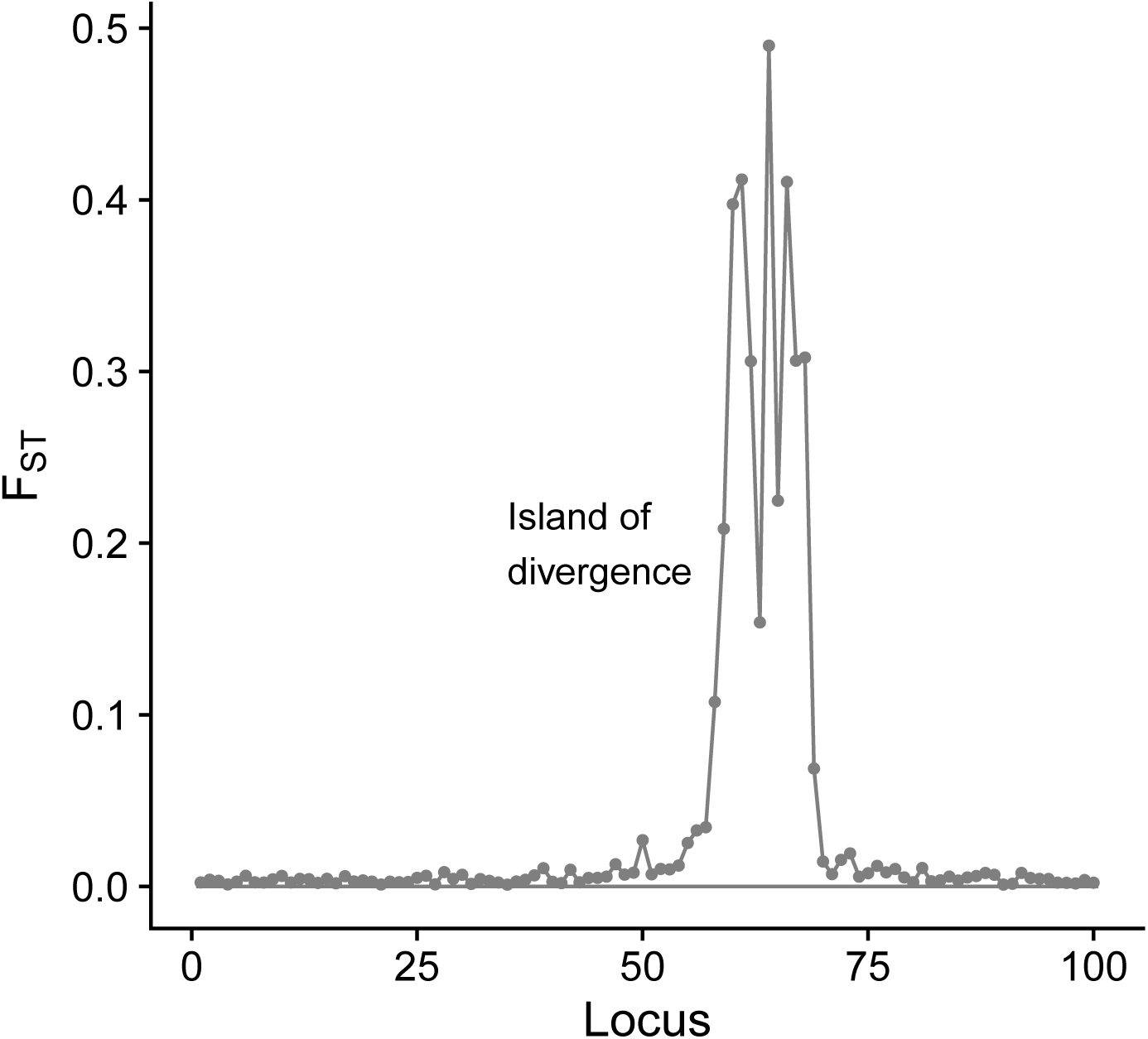
An island of divergence where many small effects at linked loci build up a bigger effect. The value of the between-habitat *F*_ST_ is shown for each of 100 loci. Around 15 loci, with higher than background *F*_ST_, are part of the island of divergence. Survival selection between habitats is given by equation (2) and the phenotype is determined as in equation (1) with *α* = 1 and *β* = 0, so there is pure genetic phenotype determination. In the model for a single trait described main text, the genetic effect *z* was determined by one diploid locus, but here we extend to the case where *z* is additively determined by many (100) loci. The additive allelic effects at each locus can vary in the interval from −0.04 to 0.04 and recombination rates between these loci are *ρ*_*zz*_ = 0.002. The migration rate is *m* = 0.06. Other parameter values: *N*_*p*_ = 200, *K* = 100, *s*_0_ = 0.1, *σ* = 1.0, *θ*_1_ = −0.75, *θ*_2_ = 0.75. The parameter values are similar to those in fig. 2.

**Figure A3:**
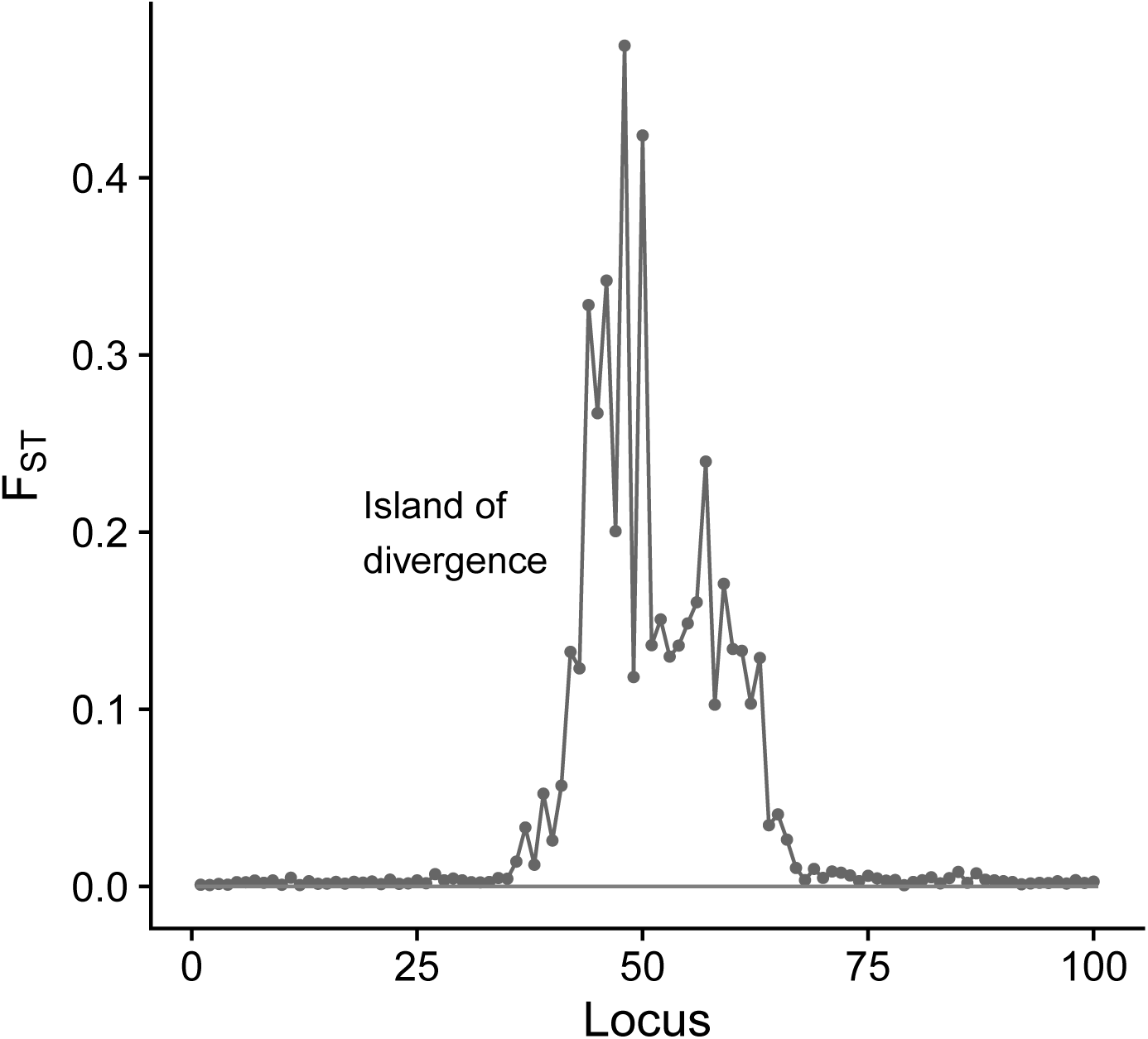
An island of divergence where many small effects at linked loci build up bigger effects for a two-trait syndrome, involving the traits *u*_1_ and *u*_2_. The value of the between-habitat *F*_ST_ is shown for each of 100 loci. Every second of these loci code for *u*_1_ and every second for *u*_2_. Around 30 loci, with higher than background *F*_ST_, are part of the island of divergence. Survival selection between habitats is given by equation (5) and the phenotype is determined as in equation (4) with *α*_1_ = *α*_2_ = 1 and *β*_1_ = *β*_2_ = 0, so there is pure genetic phenotype determination. The additive allelic effects at each locus can vary in the interval from −0.04 to 0.04 and recombination rates between loci are *ρ*_*zz*_ = 0.002. The migration rate is *m* = 0.12. The parameter values are similar to those in fig. 5.

**Author contributions**
OL designed the study, wrote and ran the computer programs, and wrote the text. SRXD, JMM, BK, and PH provided input to the design and contributed to the writing of the text.

## Acknowledgments

This work was supported by grants from the Carl Trygger Foundation (CTS 15292) to OL and by a Leverhulme Trust International Network Grant to SRXD, PH, OL, and JMM.

